# KIF1A/UNC-104 transports ATG-9 to regulate neurodevelopment and autophagy at synapses

**DOI:** 10.1101/057166

**Authors:** Andrea K. H. Stavoe, Sarah E. Hill, Daniel A. Colón-Ramos

## Abstract

Autophagy is a cellular degradation process essential for neuronal development and survival. Neurons are highly polarized cells in which autophagosome biogenesis is spatially compartmentalized. The mechanisms and physiological importance of this spatial compartmentalization of autophagy in the neuronal development of living animals are not well understood. Here we determine that, in *C. elegans* neurons, autophagosomes form near synapses and are required for neurodevelopment. We first determined, through unbiased genetic screens and systematic genetic analyses, that autophagy is required cell-autonomously for presynaptic assembly and for axon outgrowth dynamics in specific neurons. We observe autophagosomes in the axon near synapses, and this localization depends on the synaptic vesicle kinesin, KIF1A/UNC-104. KIF1A/UNC-104 coordinates localized autophagosome formation by regulating the transport of the integral membrane autophagy protein, ATG-9. Our findings indicate that autophagy is spatially regulated in neurons through the transport of ATG-9 by KIF1A/UNC-104 to regulate neurodevelopment.

## HIGHLIGHTS and eTOC Blurb

- The autophagy pathway acts cell-autonomously and in specific neurons in development
- Autophagosome biogenesis occurs in compartmentalized axonal regions near synapses
- The synaptic vesicle kinesin UNC-104/KIF1A transports ATG-9 to presynaptic sites

## INTRODUCTION

Macroautophagy (hereafter called autophagy) is an evolutionarily conserved cellular degradation process best known for its role in cellular homeostasis (Feng et al., 2014; Marino et al., 2011; Son et al., 2012; Wu et al., 2013a; Zhang and Baehrecke, 2015). While autophagy is induced under stress conditions in yeast and many mammalian cells, in neurons, autophagosome formation is a constitutively active process (Lee, 2012; Wong and Holzbaur, 2015; Xilouri and Stefanis, 2010). Basal levels of autophagy are essential for neuronal survival, and neuron-specific inhibition of the autophagy pathway results in axonal degeneration and neuronal cell death (Hara et al., 2006; Komatsu et al., 2007; Yang et al., 2013; Yue et al., 2009).

Autophagy can regulate axon morphogenesis and synaptic physiology in neurons (Binotti et al., 2015; Hernandez et al., 2012; Shehata and Inokuchi, 2014; Torres and Sulzer, 2012; Yamamoto and Yue, 2014). For example, knockdown of the autophagic protein Atg7 in murine neurons results in longer axons, while activation of the autophagy pathway with rapamycin results in shorter neurites (Ban et al., 2013; Chen et al., 2013). In *Drosophila*, disruption of autophagy decreases the size of the neuromuscular junction, while induction of autophagy increases synaptic boutons and neuronal branches (Shen and Ganetzky, 2009). These changes in the axon are dependent, at least in part, on the degradation of cytoskeletal regulatory proteins and structures (Ban et al., 2013). Consistent with these observations, autophagosomes form at the tips of actively elongating axons in cultured neurons and contain membrane and cytoskeletal components (Bunge, 1973; Hollenbeck, 1993; Hollenbeck and Bray, 1987; Maday et al., 2012).

Most of our knowledge of autophagosome biogenesis comes from studies conducted either in yeast or mammalian cell culture (Abada and Elazar, 2014; Hale et al., 2013; Reggiori and Klionsky, 2013). Less is known about how autophagy is regulated *in vivo* in multicellular organisms during development and stress (Wu et al., 2013a; Zhang and Baehrecke, 2015). Studies from *C. elegans, Drosophila* and mice have revealed that autophagy plays critical roles in the coordinated execution of developmental programs (Hale et al., 2013; Wu et al., 2013a). For example, autophagic degradation in early embryos is essential for pre-implantation development in mammals and has been linked to the degradation of maternal proteins in oocytes (Tsukamoto et al., 2008). In *C. elegans* and *Drosophila*, autophagy plays important roles in degrading paternal organelles after fertilization (Al Rawi et al., 2011; DeLuca and O’Farrell, 2012;Sato and Sato, 2011; Yang and Zhang, 2014; Zhang and Baehrecke, 2015). Similarly, P granules, structures normally present in germline cells, are degraded in somatic cells by autophagy (Lu et al., 2013; Zhang et al., 2009). In all these developmental programs, autophagy plays critical roles during development by degrading substrates at specific developmental transitions. Autophagy also plays important roles during the development of the nervous system (Boland and Nixon, 2006; Cecconi et al., 2007; Lee et al., 2013; Yamamoto and Yue, 2014). However, the specific roles of autophagy in the coordination of neurodevelopmental events are less clear.

Neurons are highly polarized cells. In primary neurons, autophagosomes are observed to form at the distal end of the axon, indicating compartmentalization and spatial regulation of autophagosome biogenesis (Ariosa and Klionsky, 2015; Ashrafi et al., 2014; Hollenbeck, 1993; Hollenbeck and Bray, 1987; Maday and Holzbaur, 2014; Maday et al., 2012; Yue, 2007). Autophagosome biogenesis requires the ordered recruitment of assembly factors to the distal axon (Maday and Holzbaur, 2014; Maday et al., 2012). How localized recruitment is regulated in neurons to specify autophagosome biogenesis is not well understood.

In this study, we conducted unbiased forward genetic screens to identify pathways involved in presynaptic assembly in *C. elegans* and identified an allele of *atg-9.* ATG-9 is known for its role in autophagosome biogenesis (Lang et al., 2000; Noda et al., 2000; Reggiori et al., 2004; Stanley et al., 2013; Wang et al., 2013; Yamamoto et al., 2012). Through genetic and cell biological approaches, we determined that the autophagy pathway is required cell-autonomously to promote presynaptic assembly in the interneuron AIY. We performed systematic analyses with cell biological markers for cytoskeletal organization, active zone position and synaptic vesicle clustering to establish that sixteen distinct autophagy genes promote proper neurodevelopment in *C. elegans.* We examined different neuron types to determine that autophagy regulates synaptic positions and axon outgrowth in specific neurons. We found that ATG-9 is transported to the tip of growing neurites or to synaptic regions by the synaptic vesicle kinesin UNC-104/KIF1A. Transport of ATG-9, in turn, regulates synaptic autophagosomes. We propose that transport of ATG-9 by UNC-104/KIF1A confers spatial regulation of autophagy in neurons.

## RESULTS

### Mutant allele *wy*56 displays defects in AIY synaptic vesicle clustering

During development, the AIY interneurons of *C. elegans* establish a stereotyped pattern of *en passant* (along the length of the axon) synaptic outputs. This pattern is reproducible across animals and displays specificity for both synaptic partners and positions (Figures 1A-1C) (Colón-Ramos et al., 2007; White et al., 1986). The positions of these *en passant* synapses in AIY are instructed by glia-derived Netrin, which directs local organization of the actin cytoskeleton, active zone localization and synaptic vesicle clustering (Colón-Ramos et al., 2007; Stavoe et al., 2012; Stavoe and Colón-Ramos, 2012). To identify the mechanisms underlying these early events that organize synaptogenesis, we performed visual forward genetic screens using the synaptic vesicle marker GFP::RAB-3 in AIY. From these screens, we identified the allele *wy56*, which displays a highly penetrant, abnormal distribution of synaptic vesicle proteins GFP::RAB-3 and SNB-1::YFP in the dorsal turn region of the AIY neurite (termed Zone 2; Figures 1E–1G and S1H).

**Figure 1.**
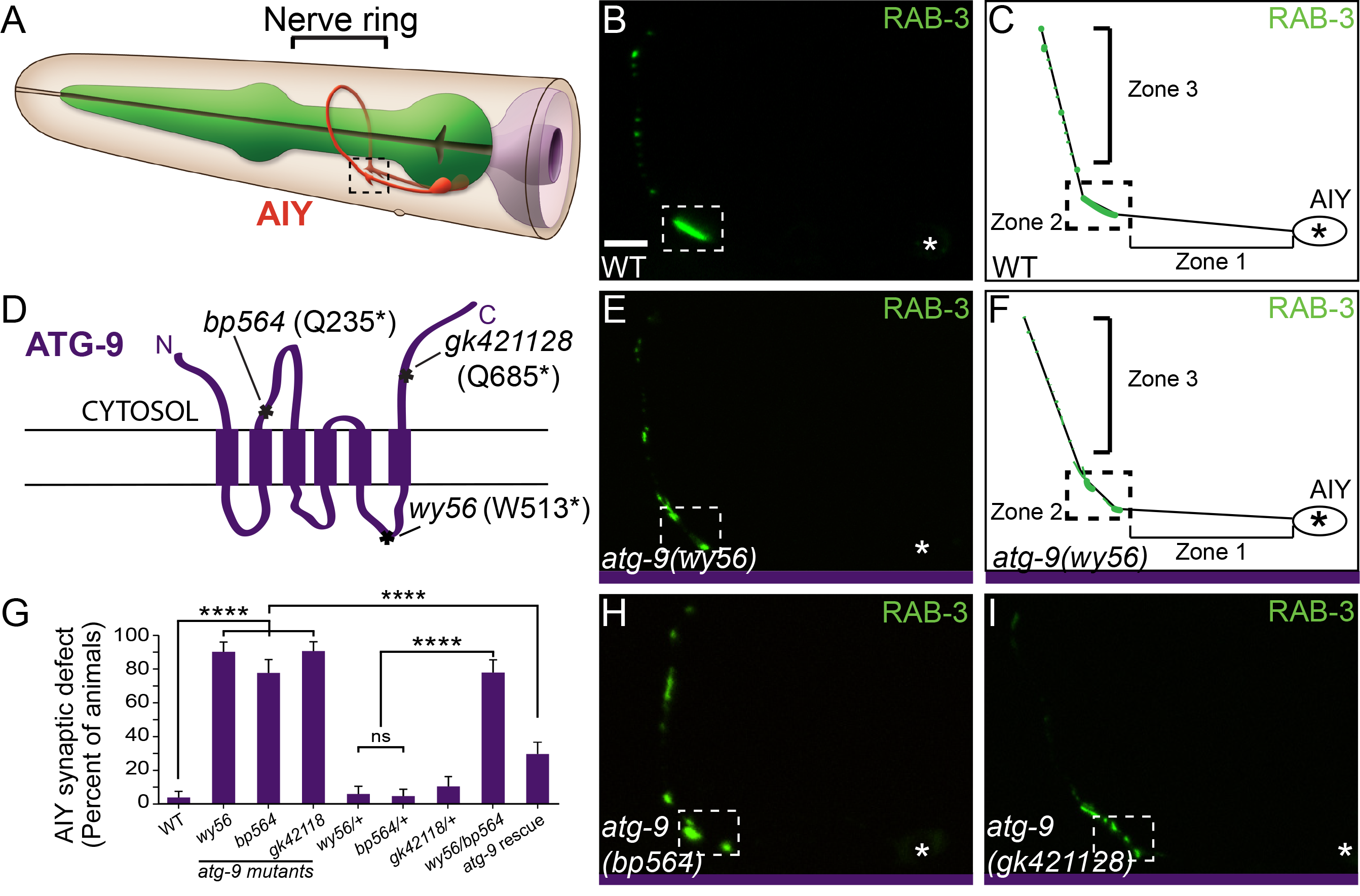
ATG-9 is required for AIY synaptic vesicle clustering. (A) Schematic of AIY interneurons (red) in the nerve ring (indicated by bracket) in the head of the worm. Reprinted with permission from wormatlas.org (Z. Altun). (B) Distribution of synaptic vesicles (visualized with GFP::RAB-3) in a representative wild type animal and represented in a schematic diagram (C). Throughout the paper, Zone 1 corresponds to an asynaptic region of the AIY neurite, Zone 2 corresponds to a synaptic rich region in the dorsal turn of the neurite and Zone 3 corresponds to a synaptic region with intermittent presynaptic clusters. We focus the characterization of our phenotypes to Zone 2, as described previously (Colón-Ramos et al., 2007; Stavoe et al., 2012; Stavoe and Colón-Ramos, 2012). (D) Schematic of ATG-9 transmembrane domains with *bp564, wy56* and *gk421128* lesions and corresponding protein effects indicated. (E) Distribution of synaptic vesicles in an *atg-9(wy56)* mutant animal and represented in a cartoon diagram (F). (G) Quantification of the AIY presynaptic phenotype in wild type, three *atg-9* mutant backgrounds, heterozygous *atg-9* animals, *atg-9 (wy56/bp564)* transheterozygotes, and *atg-9(bp464)* mutant animals that contain a pan-neuronal *(punc-14) atg-9* rescuing construct. Note that the transheterozygous animals display AIY presynaptic defects, while the single heterozygous animals are similar to wild type animals. Error bars represent 95% confidence interval. ****, P< 0.0001 between indicated groups by Fisher’s exact test. (H-I) Distribution of synaptic vesicles in *atg-9(bp564)* (H) and *atg-9(gk421128)* (I) mutant animals. Each image is a maximal projection of a confocal z-stack; the asterisk denotes the location of the cell body and the dashed box encloses AIY Zone 2. Scale bar (in B for B, E, H, and I), 5μm.

We observed that both the penetrance and expressivity of *wy56* mutant animals resembled those of other synaptogenic mutants in AIY, including mutants of actin-organizing molecules CED-5/D0CK-180, CED-10/RAC-1 and MIG-10/Lamellipodin (Stavoe et al., 2012; Stavoe and Colón-Ramos, 2012). Also similar to these actin-organizing mutants, the *wy56* defective presynaptic pattern was observed early in development, at larval stage 1 (L1), just after hatching, suggesting that the phenotype emerges during embryogenesis (Figure S2). Our findings indicate that *wy56* corresponds to a lesion in a gene required early in development for correct formation of presynaptic sites in AIY.

### *wy*56 is an allele of autophagy gene *atg-9*

To identify the genetic lesion of *wy56*, we performed single-nucleotide polymorphism (SNP) mapping, whole-genome sequencing and genetic rescue experiments. Our SNP mapping data indicate that *wy56* is located between 0.05 Mb and 0.5 Mb on chromosome V. Whole genome sequencing of *wy56* mutants revealed a point mutation in exon 8 of the *atg-9* (AuTophaGy-9) gene, resulting in a G to A nucleotide transition that converts W513 to an opal/umber stop codon (Figure 1D). Two independent alleles of *atg-9*, *atg-9*(*bp564*) and *atg-9(gk421128)*, with amber nonsense mutations at Q235 and Q685, respectively (Figure 1D) (Thompson et al.,2013; Tian et al., 2010), phenocopied the AIY presynaptic defect observed for *wy56* mutants(Figures 1H-1I). Consistent with *wy56* being an allele of *atg-9*, we also observed that *wy56* fails to complement *atg-9(bp564)* and that expression of the ATG-9 cDNA under an early panneuronal promoter *(punc-14)* rescues the *atg-9* presynaptic phenotype in AIY (Figure 1G). Together, our genetic data indicate that *wy56* is a nonsense, loss-of-function mutation in the *atg-9* gene.

### The autophagy pathway is required for clustering of synaptic vesicle proteins in AIY

The *atg-9* gene encodes a conserved, six-pass transmembrane protein that acts in the autophagy pathway (Lang et al., 2000; Noda et al., 2000; Young et al., 2006). Since ATG-9 is primarily known for regulating autophagosome biogenesis, we examined whether other components of the autophagy pathway are also required for synaptic vesicle clustering during synaptogenesis. Autophagosome biogenesis can be divided into four steps: initiation, nucleation, elongation, and retrieval. Distinct and specialized protein complexes mediate these steps, and orthologs for these protein complexes have been identified in *C. elegans* (Figures 2A, S3G and Table S1) (Melendez and Levine, 2009; Tian et al., 2010). To evaluate the requirement of each of these distinct steps in synaptic vesicle clustering, we systematically examined existing alleles for each of these orthologs using synaptic vesicle markers GFP::RAB-3 and SNB-1::YFP (Figures 2, S1 and S3).

**Figure 2.**
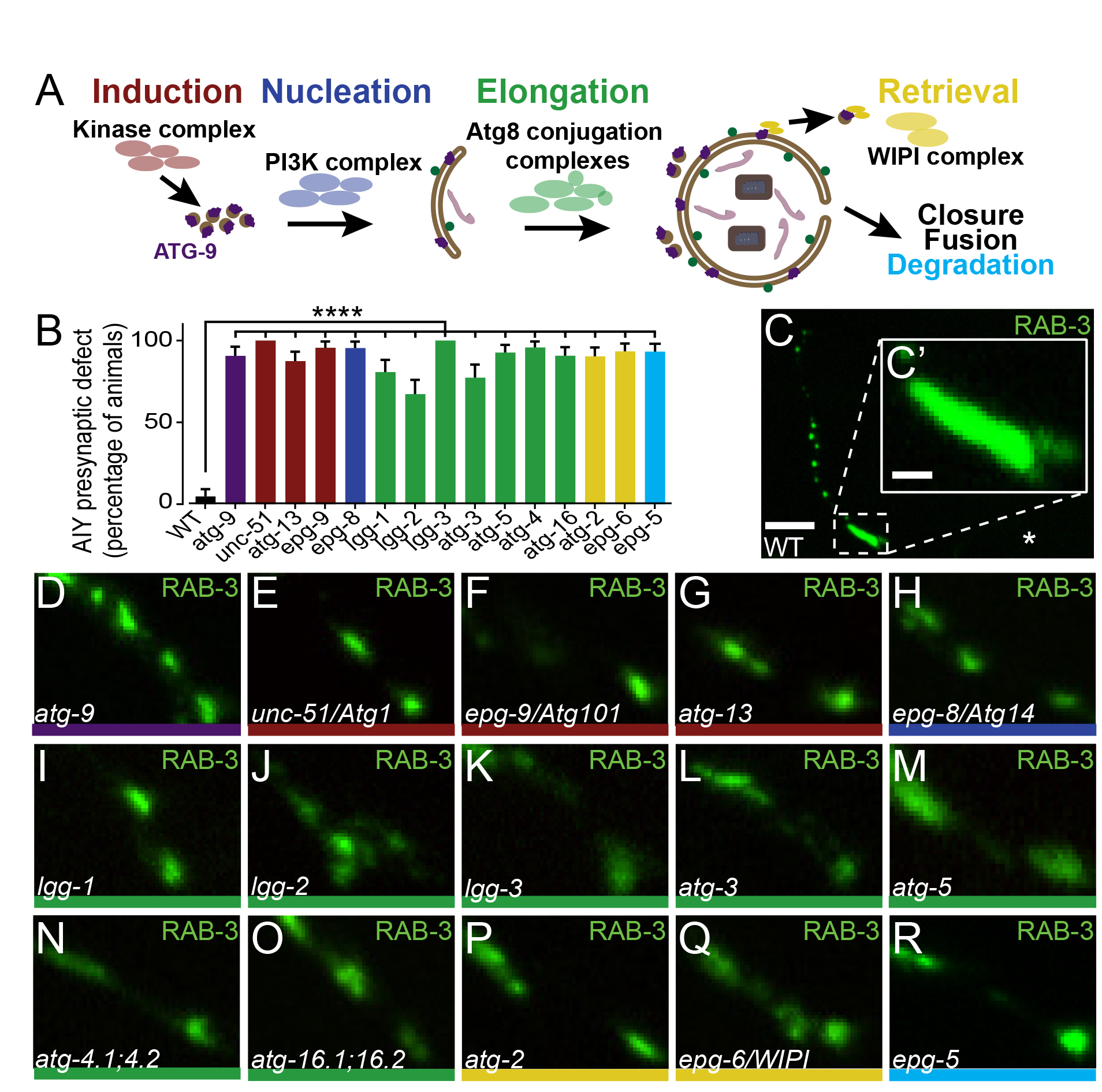
The autophagy pathway is required for AIY synaptic vesicle clustering. (A) Schematic of the autophagosome biogenesis pathway, including the general stages: induction (red), nucleation (blue), elongation (green), retrieval (yellow), closure, fusion and degradation (cyan). In all figures, colored bars under the images and in graphs denote the step of the autophagy pathway affected. (B) Quantification of the AIY presynaptic defect in wild type and autophagy pathway mutant animals. For all categories quantified, n>100 animals. Error bars represent 95% confidence interval. ****, P< 0.0001 between autophagy mutants and wild type by Fisher’s exact test. (C) Distribution of synaptic vesicles (visualized with GFP::RAB-3) in a representative wild type animal. (C’) Inset in (C) is enlarged Zone 2 region from (C). (D-R) Distribution of synaptic vesicles (visualized with GFP::RAB-3) in AIY Zone 2 of *atg-9(gk421128)* (D), *unc-51(e369)* (E), *epg-9(bp320)* (F), *atg-13(bp414)* (G), *epg-8(bp251)* (H), *lgg-1(bp500)* (I), *lgg-2(tm5755)* (J), *lgg-3(tm1642)* (K), *atg-3(bp412)* (L), *atg-5(bp484)* (M), *atg-4.1(gk127286);atg-4.2(gk430078)* double mutants (N), *atg-16.1(gk668615);atg-16.2(gk145022)* double mutants (O), *atg-2(bp576)* (P), *epg-6(bp242)* (Q), and *epg-5(tm3425)* (R) mutant animals. Each image is a maximal projection of a confocal z-stack; the asterisk denotes the location of the cell body and the dashed box encloses AIY Zone 2 in (C). Only the AIY Zone 2 region is depicted in (D-R), similar to the region depicted in (C’). Scale bar in C, 5 μm; scale bar in C’ for C’-R, 1m. See also Figures S1, S2 and S3 and Table S1

Autophagy is induced by a kinase complex composed of UNC-51/Atg1/ULK, EPG-9/Atg101 and ATG-13/EPG-1 (Feng et al., 2014; Kamada et al., 2000; Melendez and Levine, 2009; Reggiori et al., 2004). Examination of putative null alleles for *unc-51/Atg1/ULK, epg-9/Atg101* and *atg-13/epg-1* revealed highly penetrant AIY synaptic vesicle clustering defects which phenocopied those seen for *atg-9* mutant animals (Table S1). We found that 100% of *unc-51(e369)* mutant animals (n=109), 95.6% of *epg-9(bp320)* mutant animals (n=113) and 87.4% of *atg-13(bp414)* mutant animals (n=127) exhibit reduced synaptic vesicle clustering in Zone 2, as visualized with synaptic vesicle-associated markers GFP::RAB-3 and SNB-1::YFP (Figures 2B, 2E-2G, and S1B), suggesting that the initiation complex of autophagy is required for clustering synaptic vesicle proteins in AIY.

Nucleation is mediated by a PI3K complex that promotes fusion of ATG-9-containing vesicles into a phagophore (Kihara et al., 2001; Obara et al., 2006). The nucleation complex consists of LET-512/Vps34, BEC-1/Atg6, EPG-8/Atg14, and VPS-15. Most of these genes also play important roles in other essential cellular pathways, and for this reason, null mutations in these genes are unviable (Kihara et al., 2001; Obara et al., 2006; Yang and Zhang, 2011). However, we were able to examine a putative null allele for the nucleation gene *epg-8/Atg14*, which is specific to the autophagy pathway (Table S1) (Yang and Zhang, 2011). Similar to *atg-9* mutants and initiation complex mutants, *epg-8(bp251)* mutant animals exhibited highly penetrant AIY synaptic vesicle clustering defects, as visualized with synaptic vesicle markers GFP::RAB-3 and SNB-1::YFP (95.4% of *epg-8(bp251)* mutant animals, n=108; Figures 2B,2H and S1C).

Once nucleated, the isolation membrane elongates via two ubiquitin-like conjugation complexes. In *C. elegans*, these complexes include: LGG-1/GABARAP, LGG-2/LC3, LGG-3/Atg12, ATG-5, ATG-7, ATG-10, ATG-4 (ATG-4.1 and ATG-4.2 in *C. elegans)*, ATG-16 (ATG-16.1 and ATG-16.2 in *C. elegans)*, and ATG-3 (Melendez and Levine, 2009). We examined putative null alleles for *atg-5, lgg-3/Atg12, lgg-1/GABARAP* and *lgg-2/LC3*, and a hypomorphic allele of *atg-3* (Table S1). Because *atg-4* and *atg-16* both have two orthologs with redundant functions in *C. elegans* (Wu et al., 2012; Zhang et al., 2013), we built double mutant strains carrying putative null alleles for both orthologs *(atg-4.1;atg-4.2* double mutants and *atg-16.1;atg-* 16.2 double mutants; Table S1). Consistent with elongation playing an important role in synaptic vesicle clustering in AIY, we observed that all of these mutants displayed highly penetrant synaptic vesicle clustering defects in AIY as visualized by synaptic vesicle associated protein GFP::RAB-3 (all these elongation mutants display >65% penetrance, n>100 animals for all genotypes quantified; Figures 2B and 2I-2o) or SNB-1::YFP (Figure S1D).

Next, ATG-9 is retrieved from the isolation membrane by a retrieval complex before the double membrane can close, creating a mature autophagosome (Lu et al., 2011; Reggiori et al., 2004; Wang et al., 2001). Similar to our previous results, we observed that putative null alleles for retrieval genes *epg-6* (a WIPI ortholog) and *atg-2* (Table S1) displayed defects in AIY synaptic vesicle clustering, as visualized by synaptic vesicle markers GFP::RAB-3 and SNB-1::YFP (Figures 2B, 2P-2Q and S1E-S1G).

Once the autophagosome has formed, it fuses with either late endosomes or lysosomes to form the autolysosome, an organelle containing proteolytic enzymes that degrade the autophagosomal contents. In *C. elegans* and mammals, EPG-5 is required for the maturation of autolysosomes (Cullup et al., 2013; Tian et al., 2010; Zhao et al., 2013). We examined the putative null allele *epg-5(tm3425)* and observed, consistent with earlier autophagy mutants, highly penetrant AIY synaptic vesicle clustering defects as visualized with GFP::RAB-3 (93.0% of *epg-5(tm3425)* mutant animals, n=114; Figure 2B and 2R).

Together, our findings indicate that components from all stages of the autophagy pathway are required for clustering of synaptic vesicle proteins in AIY during development. Our data further suggest that ATG-9 is acting within the autophagy pathway, not independently, to direct AIY synaptic vesicle clustering during development.

### Selective autophagy genes *atg-11* and *epg-2* are not required for AIY synaptic vesicle clustering

Autophagy is best known for its nonselective role in degrading bulk cytoplasm (Mizushima and Klionsky, 2007). In neurons, nonselective autophagy locally degrades membrane and cytoskeletal components at the tips of actively elongating axons (Ban et al., 2013; Chen et al., 2013; Hollenbeck and Bray, 1987). However, the autophagy pathway can also selectively degrade specific organelles or target proteins in certain contexts. This process, known as selective autophagy, is dependent on the adaptor molecule Atg11, which interacts with cargo receptors to link specific protein targets to the autophagosome precursor membrane (Farre et al., 2013; He et al., 2006; Mao et al., 2013; Yorimitsu and Klionsky, 2005). To determine whether selective autophagy acts in AIY synaptic vesicle clustering, we examined two putative null alleles of *atg-11 (tm2508* and *(gk863004)* (Table S1). Unlike the other autophagy pathway mutants examined, we did not observe AIY presynaptic defects in *atg-11* mutants, suggesting that *atg-11*-dependent selective autophagy is not required for AIY presynaptic assembly (Figures S3B-S3C). Consistent with these findings, we also observed that putative null mutants for *epg-2* (Table S1), a proposed selective autophagy adaptor in *C. elegans* (Tian et al., 2010), also did not display AIY presynaptic defects (Figure S3D). Therefore, our data suggest that selective autophagy adaptors ATG-11 and EPG-2 are not required for AIY synaptic vesicle clustering.

### The autophagy pathway acts cell-autonomously in AIY to promote synaptic vesicle clustering

Next we examined whether autophagy acts cell-autonomously in AIY by performing mosaic analyses with representative mutants from distinct steps of the autophagy pathway. Briefly, mitotically-unstable rescuing arrays were used for each of the examined genes, and animals were scored for retention of the rescuing array in AIY and for the synaptic phenotype (Yochem and Herman, 2003). We observed that for *atg-9(bp564), lgg-1(bp500)* (elongation) and *atg-2(bp576)* (retrieval) mutants, retention of their rescuing arrays in AIY resulted in rescue of the AIY presynaptic defect, while retention of the array in other neurons (including postsynaptic partner RIA), but not in AIY, did not result in rescue of the AIY presynaptic defect (Figure 3G-3O). We also examined the endogenous expression pattern of *atg-9* and *atg-2* and observed that they are expressed in neurons, including AIY (Figure S4). Together, our data suggest that the autophagy pathway acts cell-autonomously in AIY to promote synaptic vesicle clustering.

**Figure 3.**
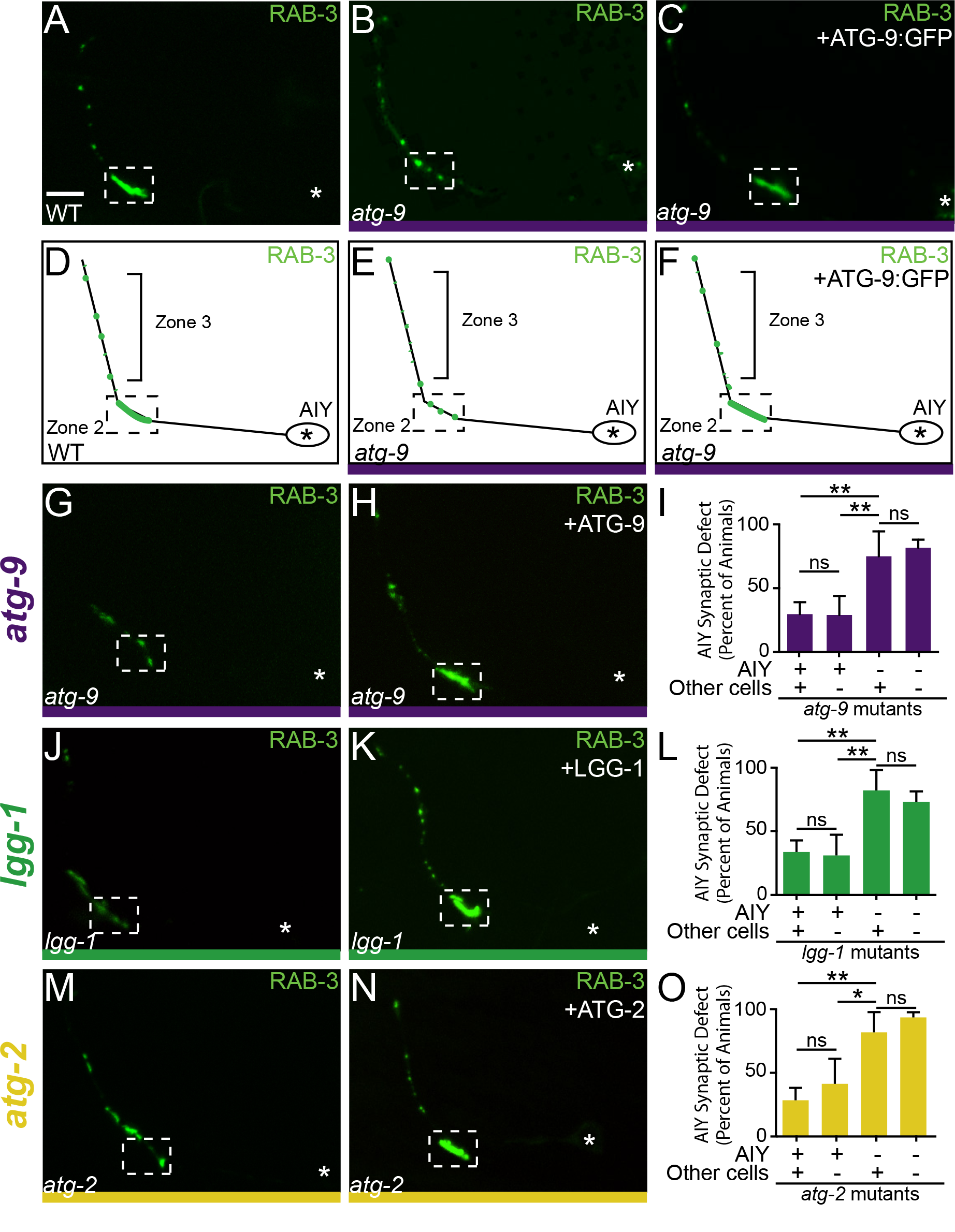
The autophagy pathway acts cell-autonomously to instruct AIY presynaptic assembly. (A) Distribution of synaptic vesicles (visualized with GFP::RAB-3) in a representative wild type animal. (B-C) Distribution of synaptic vesicles (visualized with mCh::RAB-3, pseudocolored green) in an *atg-9(bp564)* mutant animal (B) and *atg-9(bp564)* mutant expressing a panneuronal ATG-9::GFP rescuing construct (C). (D-F) Schematics of synaptic vesicle patterns in (A-C), respectively. (G-H) Distribution of synaptic vesicles in *atg-9(bp564)* mutant animals (G) and *atg-9* mutant animals expressing a rescuing array (H). (I) Quantification of rescue of *atg-9(bp564)* mutant animals expressing a rescuing array in AIY. (J-K) Distribution of synaptic vesicles in *lgg-1(bp500)* mutant animals (J) and *lgg-1* mutant animals expressing a rescuing array (K). (L) Quantification of rescue of *lgg-1* mutant animals expressing a rescuing array in AIY. (M-N) Distribution of synaptic vesicles in *atg-2(bp576)* mutant animals (M) and *atg-2* mutant animals expressing a rescuing array (N). (O) Quantification of rescue of *atg-2* mutant animals expressing a rescuing array in AIY. (I, L, O) Mutant animals expressing unstable rescuing transgenes and cell-specific cytoplasmic markers in AIY and RIA were assayed for retention of the transgene and rescue of the AIY presynaptic phenotype as described (Colon-Ramos et al., 2007). Note that retention of the rescuing transgenes in AIY, but not other cells (as determined by retention in RIA), results in rescue of the AIY presynaptic defect. Error bars represent 95% confidence interval. * *, P< 0.01; *, P< 0.05 between indicated groups by Fisher’s exact test. Each image is a maximal projection of a confocal z-stack; the asterisk denotes the location of the cell body and the dashed box encloses AIY Zone 2. Scale bar (in A for A-C, G-H, J-K, M-N), 5m. See also Figure S4.

### The autophagy pathway is required for F-actin accumulation and active zone protein localization at presynaptic sites

To understand how autophagy regulates synaptic vesicle clustering in AIY, we examined the requirement of autophagy for the different steps of presynaptic assembly. In previous studies, we determined that AIY presynaptic sites are specified through the local organization of the actin cytoskeleton and active zone proteins (Colón-Ramos et al., 2007; Stavoe et al., 2012; Stavoe and Colón-Ramos, 2012). Therefore, we examined whether the autophagy pathway was required for the localization of active zone proteins. In wild type animals, active zone protein SYD-1 colocalizes with RAB-3 in AIY presynaptic regions (Figures 4A-4D). We observed that localization of SYD-1::GFP was altered in representative autophagy mutants (Figures 4E-4L and S5). We also observed that autophagy mutants from each step of the autophagy pathway displayed significant defects in the presynaptic enrichment of the F-actin probe UtrCH::GFP (Figures 4M-4Q). Our findings indicate that autophagy acts upstream of actin organization, active zone assembly and synaptic vesicle clustering during synaptogenesis.

**Figure 4.**
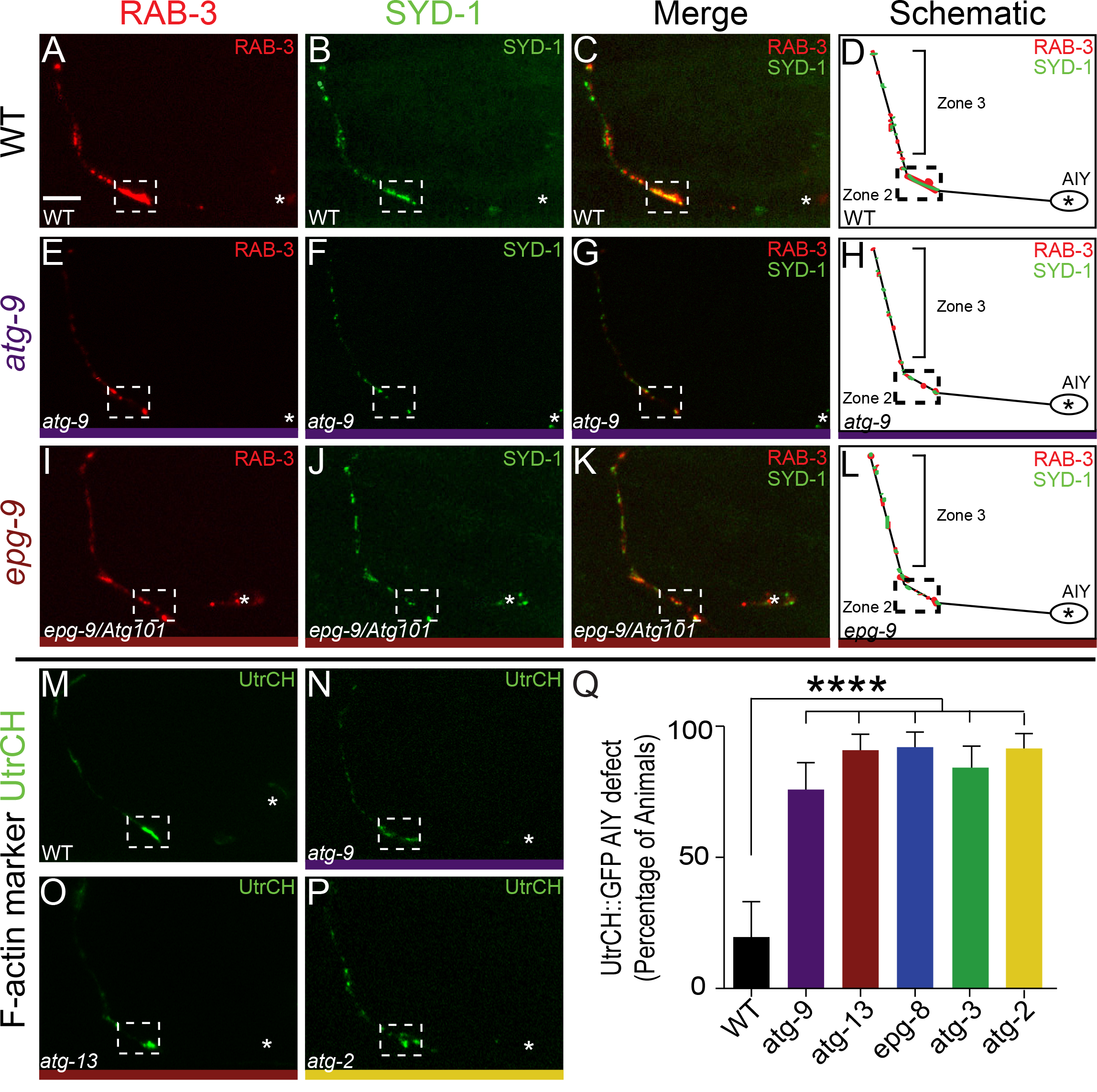
The autophagy pathway is required for active zone assembly and F-actin organization. (A-L) Distribution of synaptic vesicles in AIY (visualized with mCh::RAB-3), and localization of active zones in AIY (visualized with GFP::SYD-1) in wild type (A-C), *atg-9(bp564)* (E-G) and *epg-9(bp320)* (I-K) mutant animals. (D, H, L) Schematics of merged images (C, G, K) for the indicated autophagy mutants. (M-P) F-actin organization (visualized with UtrCH::GFP) in wild type (M), *atg-9(bp564)* (N), *atg-13(bp414)* (O), and *atg-2(bp576)* (P) mutant animals. (Q) Quantification of penetrance of F-actin Zone 2 enrichment defect in AIY. For all genotypes quantified, n>50 animals. Error bars represent 95% confidence interval. ****, P< 0.0001 between mutants and wild type by Fisher’s exact test. Each image is a maximal projection of a confocal z-stack; the asterisk denotes the location of the cell body and the dashed box encloses AIY Zone 2. Scale bar (in A for A-C, E-G, I-K, M-P), 5 μm. See also Figure S5.

### Autophagosomes accumulate in the AIY cell body and near presynaptic sites

In neurons, which are highly polarized cells, autophagosome biogenesis is spatially compartmentalized (Ariosa and Klionsky, 2015; Ashrafi et al., 2014; Maday and Holzbaur, 2014; Maday et al., 2012; Yue, 2007). Given the role of autophagy in synaptogenesis, we next examined the subcellular localization of autophagosomes in AIY by using GFP::LGG-1. LGG-1 is the *C. elegans* ortholog of Atg8/LC3, a molecule that associates with autophagosomal membranes upon autophagy induction (Alberti et al., 2010; Kabeya et al., 2000; Kirisako et al., 1999; Lang et al., 1998; Mizushima et al., 2010; Tian et al., 2010; Zhang et al., 2015). We observed that in most animals, GFP::LGG-1 was diffusely cytoplasmic throughout the neurite and localized to 2-10 puncta in the cell body of AIY (data not shown). In 33% of wild type animals (n=123), we observed LGG-1 puncta in the AIY neurite (Figures 5A, 5D and 5M). The LGG-1 puncta were most frequently located in the synapse-rich region Zone 2 and the region proximal to the cell body (Zone 1). This subcellular localization was not observed when we expressed GFP::LGG-1(G116A) (Figures 5B,5E and 5M), which contains a point mutation that prevents LGG-1 lipidation and conjugation to autophagosomes (Mizushima et al., 2010; Zhang et al., 2015).

To better understand the subcellular localization of autophagosomes in AIY, we examined GFP::LGG-1 in autophagy mutant backgrounds. LGG-1 fails to become conjugated to the autophagosome membrane in *atg-3* mutants (Ichimura et al., 2000; Tanida et al., 2002; Tian et al., 2010). We observed that fewer GFP::LGG-1 puncta formed in *atg-3(bp412)* mutants, both in the AIY cell body and in the neurites, consistent with a reduction of LGG-1 association with autophagosomes in these mutants (n=103; Figures 5C,5F and 5M). We then examined *atg-2(bp576), epg-6(bp424)* and *epg-5(tm3425)* mutants, all lesions in genes for late steps of autophagosome biogenesis and all known to result in the accumulation of defective autophagosomes (Lu et al., 2011; Mizushima et al., 2010; Shintani et al., 2001; Tian et al., 2010; Zhang et al., 2015). As expected, we observed a higher penetrance of animals displaying puncta in AIY neurites when we blocked these late steps of the autophagy pathway (54% of *atg-2(bp576)* mutants, 59% of *epg-6(bp424)* mutants and 55% of *epg-5(tm3425)* mutants; Figures 5G-5M). Together, our results indicate that autophagosomes accumulate in the AIY cell body and near presynaptic sites and suggest that autophagosome biogenesis is spatially compartmentalized in AIY. Our findings are consistent with studies conducted in cultured neurons, which revealed that actively elongating axons contain autophagosomes (Bunge, 1973; Hollenbeck, 1993; Hollenbeck and Bray, 1987; Maday et al., 2012). In these studies, it was hypothesized that local autophagy is critical for degrading substrates at axonal subcellular structures (Hollenbeck, 1993; Hollenbeck and Bray, 1987). How local autophagy is regulated to confer spatial selectivity during substrate degradation is not well understood.

**Figure 5.**
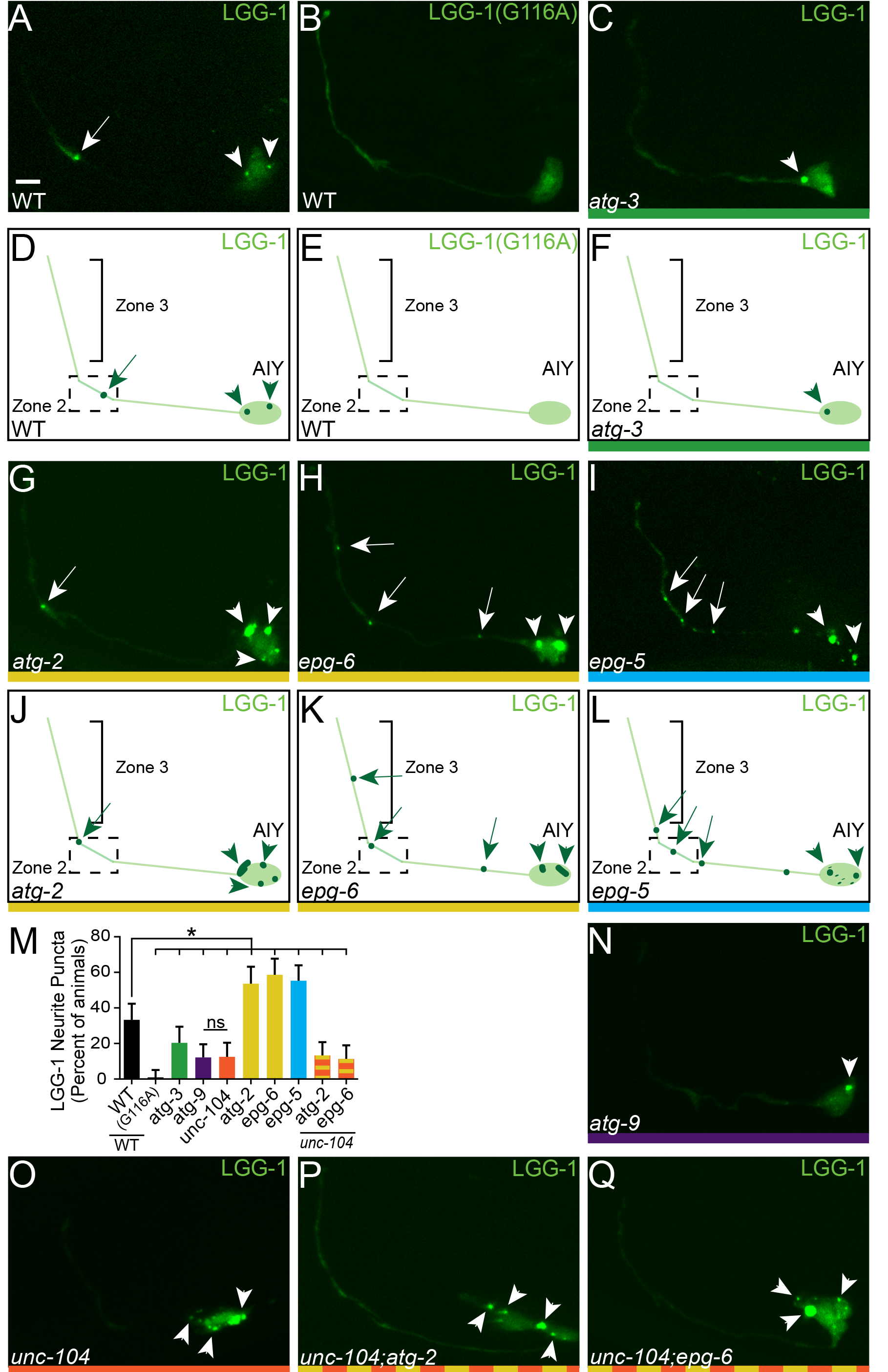
Autophagosomes are present in AIY synaptic regions. (A) Distribution of autophagosomes (visualized with GFP::LGG-1) in AIY in a wild type animal. (B) Visualization of GFP::LGG-1(G116A) in AIY in a wild type animal. LGG-1(G116A) is a point mutant incapable of associating with autophagosomes, used here as a control (Mizushima et al., 2010; Zhang et al., 2015). Note the absence of GFP::LGG-1 puncta in the neurite and cell body. (C) Distribution of autophagosomes in an *atg-3(bp412)* mutant animal. (D-F) Schematics of GFP::LGG-1 distribution throughout AIY in A-C, respectively. (G-I) Distribution of autophagosomes in *atg-2(bp576)* (G), *epg-6(bp242)* (H) and *epg-5(tm3425)* (I) mutant animals with respective schematics in (J-L). (M) Quantification of the percentage of animals containing LGG-1 puncta in the neurite of AIY in wild type and mutant animals. For all categories quantified, n>100 animals. Error bars represent 95% confidence interval. *, P< 0.05 between mutants and wild type by Fisher’s exact test. (N-Q) Distribution of autophagosomes in *atg-9(bp564)* (N), *unc-104(e1265)* (O), *unc-104(e1265);atg-2(bp576)* (P), and *unc-104(e1265);epg-6(bp242)* (Q) mutant animals. Each image is a maximal projection of a confocal z-stack; arrows denote the location of LGG-1 puncta in the neurite and arrowheads denote the location of LGG-1 puncta in the cell body. Scale bar (in A for A-C, G-I, and N-Q), 5 μm.

### ATG-9 localizes to AIY presynaptic regions in an UNC-104/KIF1A-dependent manner

To determine how the location of autophagy is specified, we examined the subcellular localization of ATG-9, the only multipass transmembrane protein that is part of the core autophagy pathway (Feng et al., 2014; Noda et al., 2000; Young et al., 2006). In the nerve ring, we observed that the subcellular localization of a rescuing ATG-9::GFP transgene was reminiscent to that of panneuronally-expressed synaptic vesicle-associated protein, RAB-3 (Figures 6A,6C-6E) (Mahoney et al., 2006). We then examined the endogenous localization of ATG-9 by creating transgenic animals in which the genomic ATG-9 locus was modified with a CRISPR-based knock-in of ATG-9::GFP (Figure 6B). Consistent with ATG-9 localizing to synaptic sites, we observed that the endogenous localization of ATG-9 was similar to panneuronally-expressed GFP::RAB-3 and ATG-9::GFP (Figure 6C-6G). To better understand the subcellular localization of ATG-9, we expressed ATG-9::GFP cell-specifically in AIY. Consistent with the endogenous expression pattern, we also observed that ATG-9 was enriched at presynaptic regions and colocalized with RAB-3 in AIY (Figure 6I-6K). Our findings indicate that ATG-9 localizes to presynaptic sites in neurons.

**Figure 6.**
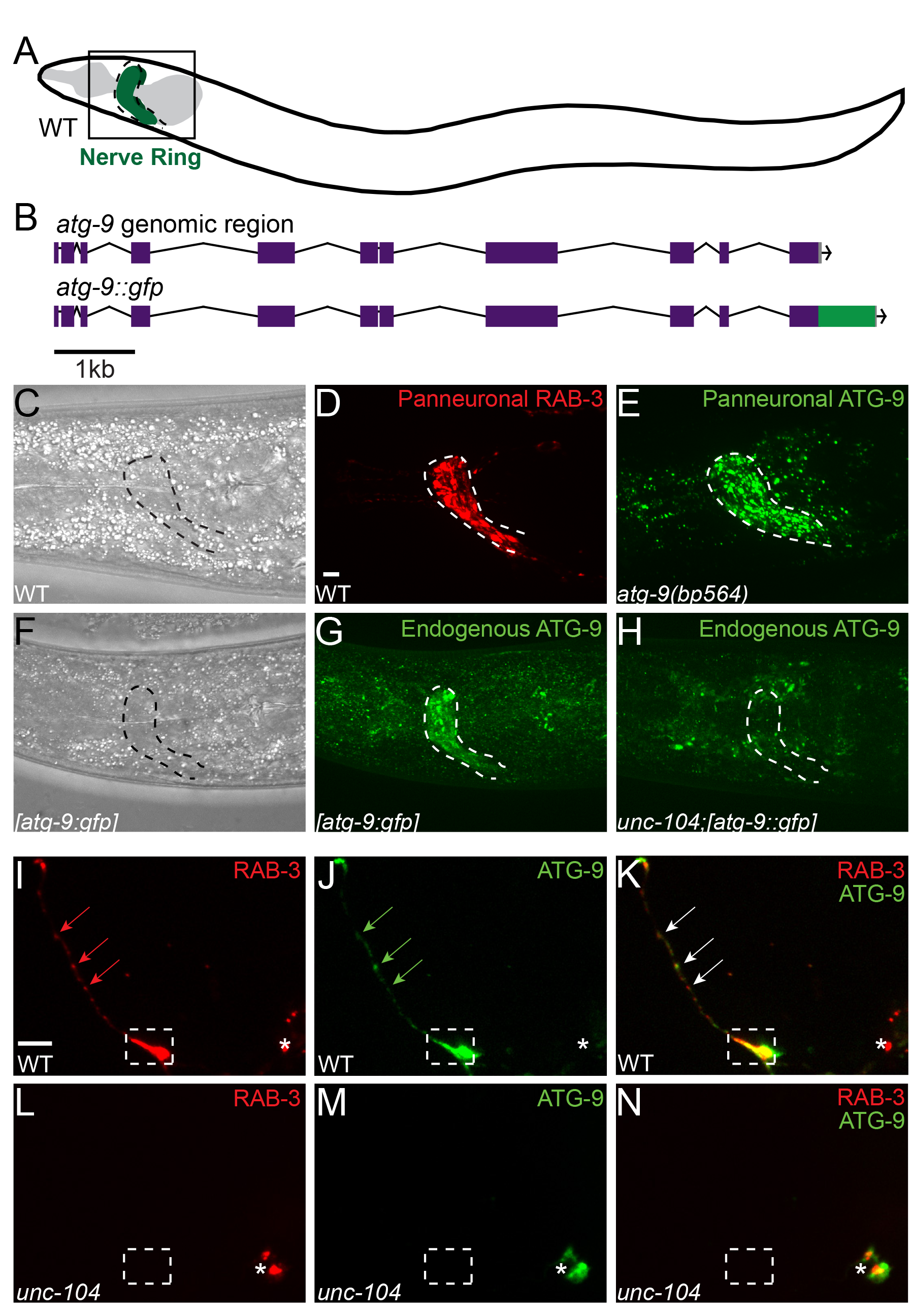
ATG-9 transport to AIY presynaptic zones requires KIF1A/UNC-104. (A) Schematic of the nematode nerve ring (green, encircled by a dotted line) in *C. elegans.* (B) Schematic of the *atg-9* genomic region in a wild type genome (top) and in the CRISPR knock-in with the *egfp* coding sequence inserted in-frame after *atg-9* (bottom). (C-D) Location of the nerve ring, referenced with a transmitted light image (C, position outlined with a dotted line) and visualized with GFP::RAB-3 (pseudocolored red) expressed panneuronally with *Prab-3* in a wild type animal (D). (E) ATG-9::GFP expressed panneuronally with *Punc-14.* (F-H) Location of the nerve ring referenced with a transmitted light image (F, position outlined with a dotted line) and distribution of endogenous ATG-9 in the head of a wild type animal, visualized with CRISPR insertion of eGFP at the C-terminus of the genomic *atg-9* in wild type animals (G) and *unc-104(e1265)* mutant animals (H). Note that ATG-9 is no longer enriched in the nerve ring in *unc-104* mutants. (I-N) Distribution of synaptic vesicles (visualized with mCh::RAB-3) (I and L) and ATG-9::GFP (J and M) in AIY (merge in K and N) in wild type (I-K) and *unc-104(e1265)* mutant animals (L-N). Note that RAB-3 and ATG-9 colocalize in AIY Zones 2 and 3 in wild type animals (dashed box and arrows) and that localization of both the RAB-3 and ATG-9 proteins is restricted to the cell body in *unc-104* mutants in AIY (L-N). Each image is a maximal projection of a confocal z-stack; in I-N, the asterisk denotes the location of the AIY cell body; the dashed box encloses AIY Zone 2. Scale bar (in D for D-E, G-H and in I for I-N), 5 μm.

ATG-9 is required for autophagosome biogenesis (He et al., 2009; Lang et al., 2000; Orsi et al., 2012; Reggiori and Tooze, 2012; Wang et al., 2013; Yamamoto et al., 2012; Young et al., 2006). In yeast, Atg9 localizes to small vesicles that nucleate to form the phagophore (Suzuki et al., 2015; Wang et al., 2013; Yamamoto et al., 2012). To understand how ATG-9 localizes to synaptic regions, we examined its subcellular localization in mutants for kinesins implicated in neuronal transport. UNC-14, a kinesin-1/KIF5 adaptor for neuronal transport of vesicles and synaptic precursors, interacts with UNC-51/ATG-1 to regulate axon outgrowth and neurodevelopment (Abe et al., 2009; Brown et al., 2009; Lai and Garriga, 2004; Ogura et al., 1997; Sakamoto et al., 2005). We did not observe any defects in synaptic vesicle clustering or ATG-9 localization in *unc-14(e57)* mutants (data not shown). Similarly, *unc-116(e2310)* mutants lacking the kinesin-1/KIF5 heavy chain and *unc-16(ju146)* mutants lacking kinesin-1/KIF5 regulator JIP3/Sunday Driver also did not display defects in ATG-9 localization (data not shown) (Byrd et al., 2001; Patel et al., 1993; Sakamoto et al., 2005; Yang et al., 2005).

However, examination of the synaptic vesicle kinesin UNC-104/KIF1A revealed a requirement for this kinesin in ATG-9 transport. In *unc-104(e1265)* mutant animals, synaptic vesicles fail to be transported to synapses and are instead restricted to the cell-body (Hall and Hedgecock, 1991; Otsuka et al., 1991). Interestingly, we observed that in *unc-104(e1265)* mutant animals, ATG-9 does not localize to synaptic regions and that its localization is restricted to the cell body (Figures 6H and 6L-6N). Our findings indicate that ATG-9 localizes to presynaptic sites in an UNC-104/KIF1A-dependent manner.

### Accumulation of autophagosomes at synaptic regions is dependent on UNC-104/KIF1A

UNC-104/KIF1A is a neuron-specific motor protein that regulates anterograde axonal transport of synaptic vesicle precursors (Otsuka et al., 1991). UNC-104/KIF1A-regulated distribution of synaptic vesicles and active zone proteins controls the spatial distribution of synapses during development (Wu et al., 2013b). Therefore, by regulating the transport of synaptic cargo to precise sites during development, the kinesin UNC-104/KIF1A provides spatial specificity for synaptogenesis.

Given the observed role of UNC-104/KIF1A in transporting ATG-9 to presynaptic sites and the known role of ATG-9 in autophagosome biogenesis, we hypothesized that localization of ATG-9 by UNC-104/KIF1A would regulate the spatial distribution of autophagosomes to presynaptic sites. To test this hypothesis, we first observed autophagosomes in *atg-9* mutant animals. Consistent with our hypothesis, we detected a significant reduction in the number of *atg-9(bp564)* mutant animals with autophagosomes in presynaptic regions (Figures 5M-5N). Next we examined the requirement of UNC-104/KIF1A for autophagosome enrichment in synaptic regions by visualizing GFP::LGG-1 in *unc-104(e1265)* mutants. We observed that the number of animals with GFP::LGG-1 puncta in the AIY neurites was significantly reduced in *unc-104(e1265)* mutants and that *unc-104(e1265)* animals phenocopied *atg-9(bp564)* mutants in terms of the penetrance of the phenotype (Figures 5M and 5O).

If UNC-104/KIF1A is important for local formation of autophagosomes, then it should act upstream of late autophagy genes *atg-2 and epg-6* to suppress the accumulation of GFP::LGG-1 puncta seen in these mutants. Indeed, we observed that *atg-2(bp576);unc-104(e1265)* and *epg-6(bp424);unc-104(e1265)* double mutants exhibited a significant reduction in the accumulation of autophagosomes in AIY presynaptic regions compared to *atg-2(bp576)* and *epg-6(bp424)* single mutants, indicating that *unc-104* is epistatic to *epg-6* and *atg-2* (Figures 5M, 5P and 5Q). Together our findings indicate that the presence of autophagosomes in synaptic regions is dependent on UNC-104/KIF1A and suggest that UNC-104/KIF1A regulates the spatial distribution of autophagosomes to presynaptic sites by mediating transport of ATG-9.

### Autophagy is required in PVD for axon outgrowth

To better understand the role of autophagy in neurodevelopment, we examined multiple neuron classes in autophagy mutants for defects in neurodevelopmental events, including axon outgrowth, axon guidance and synaptic positioning. Interestingly, most of the neurons examined (HSN, RIA, DA9, RIB, and NSM) did not display phenotypes in the examined categories (Figures S6A-S6F and data not shown). In the nociceptive sensory neuron PVD (Figure 7A), we observed that autophagy mutants were required for the length of the PVD axon at larval stage 4 (L4), but not for the morphology or timing of dendritic branching (Figures 7A-7G and data not shown).

**Figure 7.**
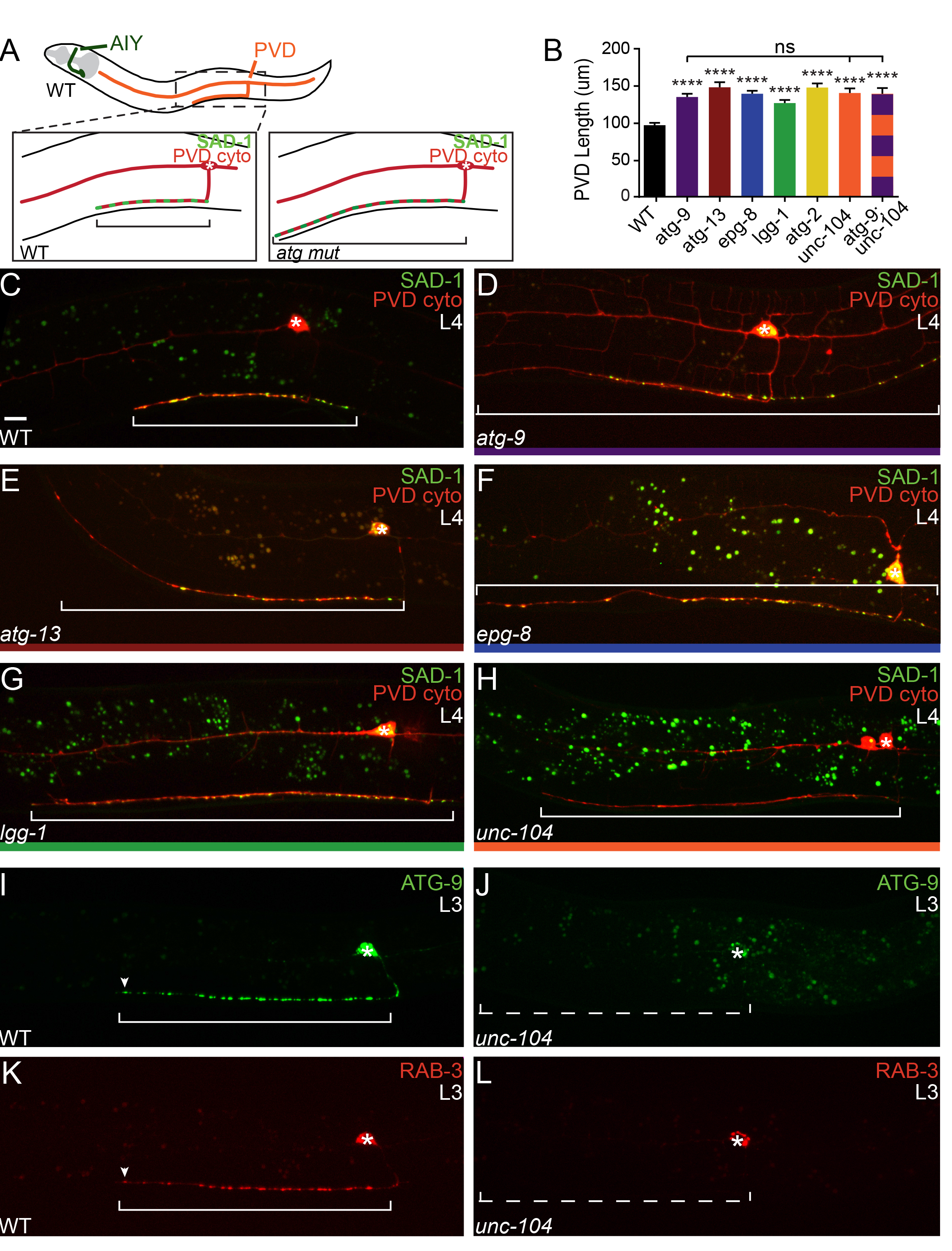
The autophagy pathway regulates the rate of PVD axon outgrowth. (A) Diagram of the morphology of AIY (green) and PVD (orange) neurons in *C. elegans.* Schematics of PVD axon morphology (red) and presynaptic sites (green) in wild type (left box) and autophagy mutant animals (right box). (B) Quantification of PVD axon length in wild type
and autophagy mutant L4 animals. Note that PVD axon length is longer in autophagy and *unc-104* mutant animals compared to wild type. Error bars represent standard error of the mean. ****, P< 0.0001 between mutants and wild type by One-way ANOVA (Tukey’s *post-hoc* analysis). (C-H) PVD morphology (visualized with cytoplasmic mCh) and presynapses (visualized with SAD-1::GFP) in the axon of PVD in wild type (C), *atg-9(bp564)* (D), *atg-13(bp414)* (E), *epg-8(bp251)* (F), *lgg-1(bp500)* (G), and *unc-104(e1265)* (H) mutant L4 animals. (I-L) Distribution of ATG-9::GFP (I and J) and synaptic vesicles (visualized with mCh::RAB-3) (K and L) in PVD in wild type (I and K) and *unc-104(e1265)* mutant animals (J and L). Note that RAB-3 and ATG-9 colocalize at presynaptic sites and at the tip of the axon in wild type animals (arrowheads in I and K) and that localization of both markers is restricted to the cell body in *unc-104* mutants (J and L). Each image is a maximal projection of a confocal z-stack; the asterisk denotes the location of the cell body; the bracket denotes length of the PVD axon; and the dotted bracket (J and L) denotes the PVD axon (not visible in the image due to absence of presynaptic protein localization). Scale bar (in C for C-L), 5 μm. See also Figure S6.

The PVD axon begins to grow during larval stage 2 (L2) and continues to grow along the ventral nerve cord into adulthood (Maniar et al., 2012; Smith et al., 2010). To understand the requirement of autophagy in PVD axon outgrowth, we also measured PVD axon length in L3 and adult stages in wild type and mutant animals. We observed that the rate of PVD axon outgrowth in mutants was significantly higher than the rate observed in wild type animals (Figures 7B and S6G). These data demonstrate that autophagy is required to regulate the rate of PVD axon outgrowth during development. Our findings also indicate that autophagy is required in individual neurons to regulate distinct and specific neurodevelopmental events *in vivo.*

### UNC-104/KIF1A is required for ATG-9 localization and axon outgrowth in PVD

We next examined the localization of ATG-9 in PVD. Similar to AIY, we observed that ATG-9::GFP localizes in a punctate pattern in the PVD axon (Figure 7I) and colocalizes with the synaptic vesicle marker mCh::RAB-3 (Figure 7K). Interestingly, we observed that in actively elongating neurons, both ATG-9::GFP and mCh::RAB-3 localize to the growing tip of the PVD axon (Figure 7I and 7K; arrowheads). In *unc-104(e1265)* mutant animals, both ATG-9 and RAB-3 did not localize to presynaptic sites or to the tip of the PVD axon and were observed instead in the PVD cell body (Figure 7J and 7L). These data suggest that, in PVD, as in AIY, UNC-104/KIF1A is important for the transport of ATG-9 in the axon.

In PVD, *atg-9* mutants, like other autophagy mutants, display longer axons than wild type animals (Figure 7B). We hypothesized that if UNC-104/KIF1A was required in PVD for ATG-9 localization and autophagy, *unc-104* mutants would phenocopy *atg-9* mutant animals, with longer PVD axons. Consistent with our hypothesis, we observed that *unc-104(e1265)* mutants phenocopied, both in terms of penetrance and expressivity, the *atg-9* mutant PVD axon length phenotype (Figures 7B and 7H). To our knowledge, this is the first time an axon outgrowth phenotype has been noted for the synaptic vesicle kinesin UNC-104/KIF1A. We hypothesized that if this newfound phenotype emerged from defects in ATG-9 transport, then *unc-104(e1265);atg-9(bp564)* double mutants would not enhance the rate of outgrowth or the length of either single mutant. Indeed, genetic analyses of *unc-104(e1265);atg-9(bp564)* double mutant animals revealed no enhancement of the PVD axon length of either single mutant, consistent with our model that *unc-104* and *atg-9* act in the same pathway to regulate PVD axon outgrowth (Figure 7B). Together our findings suggest that UNC-104/KIF-1A is important for ATG-9 transport, which in turn regulates autophagy and neurodevelopmental events.

## DISCUSSION

Autophagy regulates specific neurodevelopmental events *in vivo*. In cultured neurons, autophagosomes are observed at the tips of actively elongating axons (Bunge, 1973; Hollenbeck, 1993; Hollenbeck and Bray, 1987; Maday and Holzbaur, 2014; Maday et al., 2012). However, less is known about the specific roles of autophagy in the execution of neurodevelopmental programs, especially in the context of intact, living animals. Our systematic, *in vivo* and single-cell analyses of the role of autophagy during neurodevelopment indicate that autophagy is required for distinct and specific stages of neurodevelopment in different neuron types of *C. elegans.* We observe that in the interneuron AIY, autophagy is required to regulate presynaptic assembly, and 16 distinct autophagy mutants display consistent and specific defects in synaptogenesis. In a different neuron (sensory neuron PVD), autophagy is specifically required to regulate the rate of axon outgrowth. However, autophagy is not required for the neurodevelopment of all neurons, as several neuron classes did not reveal phenotypes in axon outgrowth, guidance or presynaptic positions in the autophagy mutants. Therefore, *in vivo*, autophagy plays precise and cell-specific roles during development to contribute to the formation of the nervous system.

Our cell biological and genetic evidence suggest that autophagy controls neurodevelopment by directly or indirectly regulating cytoskeletal structures in the axon. During development, neuronal structures such as synapses and growth cones are dynamically formed, altered or eliminated in response to developmental cues (Kolodkin and Tessier-Lavigne, 2011). Underlying the transitions during neurodevelopment are mechanisms that regulate cytoskeletal dynamics (Nelson et al., 2013; Shen and Cowan, 2010). In our study, we found that autophagy mutants phenocopied previously identified mutations in genes that regulate the actin cytoskeleton during presynaptic assembly (Colón-Ramos et al., 2007; Stavoe et al., 2012; Stavoe and Colón-Ramos, 2012). We also found that disruption of the autophagy pathway in AIY results in disordered cytoskeletal structures, abnormal active zones and mislocalized synaptic vesicles. Our findings are consistent with studies in *Drosophila* that demonstrate that autophagy regulates the development of neuromuscular junctions and with studies in vertebrates that demonstrate that autophagy-dependent changes in axon length rely on the degradation of cytoskeletal regulatory proteins (Ban et al., 2013; Shen and Ganetzky, 2009). Our findings are also consistent with studies that demonstrate that, in actively elongating axons, autophagosomes form at the tip of the axons and contain cytoskeletal components (Hollenbeck, 1993; Hollenbeck and Bray, 1987; Maday et al., 2012) and with studies that showed that induction of autophagy results in degradation of cytoskeletal components and inhibition of neurite outgrowth (Chen et al., 2013). However, loss of autophagy in neurons does not result in the pleiotropic defects one would expect from general loss of cytoskeletal regulation or cellular homeostasis. Instead, autophagy mutants display precise neurodevelopmental phenotypes in specific neurons, suggesting that regulation of autophagy is required during development. In yeast and mammalian cells, autophagosome size and number are tightly controlled through transcriptional and post-translational mechanisms (Jin and Klionsky, 2014a, Jin and Klionsky, 2014b). We hypothesize that similar mechanisms might regulate the precise deployment of autophagosomes in metazoan neurons, thereby modulating controlled degradation of structures during neurodevelopmental transitions.

Our study demonstrates that autophagosomes are compartmentalized in the axons of living animals. Previous studies have demonstrated that in cultured neurons, autophagosome biogenesis occurs in the distal axon, suggesting regulated segregation of autophagosome biogenesis in neurons (Bunge, 1973; Hollenbeck, 1993; Hollenbeck and Bray, 1987; Maday and Holzbaur, 2014; Maday et al., 2012). We now show that compartmentalization of autophagosomes in neurons is also observed *in vivo* and that autophagosomes are enriched in synaptic regions. This synaptic enrichment might confer a regulatory step by localizing the spatial activity of this cellular degradation pathway. We find that compartmentalization of autophagosomes is regulated by the local transport of ATG-9-containing vesicles. Most autophagy proteins are cytosolic, and their association with the autophagosome can be induced through post-translational modifications (Xie et al., 2015). However, ATG-9 is a six-pass transmembrane protein and the only integral membrane protein that is part of the core machinery of the autophagy pathway (Lang et al., 2000; Noda et al., 2000; Young et al., 2006).In yeast, Atg9 localizes to small (30-60nm) vesicles and promotes the formation of the autophagosome precursor (or isolation) membrane (Yamamoto et al., 2012). Little is known regarding the regulated transport of these Atg9-containing vesicles, or if their transport limits the sites of autophagosome biogenesis. In our study, we observed that ATG-9 localizes to presynaptic regions and to the tip of the growing axon. Our findings in *C. elegans* neurons are consistent with studies in mammalian neurons, which demonstrate that Atg9 is enriched in varicosities in axons and colocalizes with synaptic proteins (Tamura et al., 2010). We determined that the localization of ATG-9 to the axon is regulated by the synaptic vesicle kinesin UNC-104/KIF1A and that disruption of ATG-9 transport in *unc-104* mutants resulted in a reduced number of animals with autophagosomes in AIY neurites. Our findings provide mechanistic insights on how transport of the integral membrane protein ATG-9 instructs the spatial specificity of autophagosome biogenesis to presynaptic compartments and the distal axon.

Taken together, we propose the following model to explain how local regulation of autophagy during development could both restrain axon outgrowth in PVD and promote presynaptic assembly in AIY. We hypothesize that the UNC-104/KIF1A-dependent delivery of ATG-9 to the PVD growth cone activates autophagy and remodels the growth cone, potentially through degradation of growth cone components. This results in slower growth cone velocity, while loss of autophagy results in unrestricted axon outgrowth in PVD. This model is consistent with findings from cell culture and mammalian neurons, which reveal that disruption of autophagy results in longer neurites, while promotion of autophagy results in shorter neurites; these studies also reveal that autophagosomes form at the tips of actively elongating axons and contain cytoskeletal components (Ban et al., 2013; Bunge, 1973; Chen et al., 2013; Hollenbeck, 1993; Hollenbeck and Bray, 1987; Maday and Holzbaur, 2014; Maday et al., 2012). In AIY, formation of the synaptic-rich region Zone 2 is partially instructed by the extracellular cue Netrin (Colón-Ramos et al., 2007). We hypothesize that autophagy may be important for locally remodeling subcellular structures, such as the actin cytoskeleton, in response to this developmental cue to facilitate presynaptic assembly.

In summary, our findings suggest that in neurons of living animals, regulated transport of autophagy components, such as ATG-9, permits compartmentalized autophagosome biogenesis and progression of neurodevelopmental events. Autophagy has also been implicated in postdevelopmental events in neurons, such as synaptic transmission and vesicle recycling (Binotti et al., 2015; Hernandez et al., 2012; Wang et al., 2015). Although we focused on the characterization of developmental phenotypes, we note that autophagy protein expression persists in adults and hypothesize that the mechanisms reported here could influence synaptic physiology and function postdevelopmentally.

## EXPERIMENTAL PROCEDURES

### Strains and genetics

All worms were cultivated at 20°C on NGM plates seeded with OP50 *E. coli.* As a wild type reference strain, we utilized N2 Bristol worms. We obtained the following mutant strains from the *Caenorhabditis* Genetics Center:*unc-51/atg-1(e369)*, *atg-4.1*(*gk127286*), *atg-4.2*(*gk430078*), *atg-6/bec-1*(*ok691*), *atg-6/bec-1*(*gk202351*), *atg-9*(*gk421128*), *atg-11/epg-7*(*gk863004*), *atg-16.1*(*gk668615*), *atg-16.2*(*gk145022*), *atg-18*(*gk447069*), *epg-2*(*gk104842*), *unc-104*(*e1265*), and NM2415 (jsIs682 [Prab-3::gfp::rab-3; pJM23]). We also obtained alleles from the Hong Zhang laboratory at the Institute of Biophysics, Chinese Academy of Sciences: *atg-2*(*bp576*), *atg-3*(*bp412*), *atg-5*(*bp484*), *atg-7*(*bp422*), *lgg-1*(*bp500*), *atg-9*(*bp564*), *atg-13*/*epg-1*(*bp414*), *epg-8*/*atg14(bp251)*, *epg-6*/*WIPI*(*bp242*), and *epg-9/Atg101(bp320).* From the Mitani laboratory at the Tokyo Women’s Medical University School of Medicine, we obtained: *lgg-2(tm5755)*, *atg-11/epg-7(tm2508)*, *lgg-3/atg12(tm1642)*, *epg-5(tm3425)*, and *atg-7(tm2976)*. We obtained CX9797 (kyIs445 [Pdes-2::mCh::rab-3; Pdes-2::sad-1::gfp]) from the Bargmann lab at the Rockefeller University, NC1686 (wdIs51 [Pf49h12.4::gfp, unc-119 rescue] from the Miller lab at Vanderbilt University and *wyIs45* [Pttx-3::gfp::rab-3], *wyIs93* [Pglr-3::glr-1::gfp; Pglr-3::mCh::rab-3] and *wyIs97* [Punc-86::myrgfp; Punc-86::mCh::rab-3] from the Shen lab at Stanford University. Specific allele lesion information can be found in Table S1.

### Molecular biology and transgenic lines

We created pSM vector-derived plasmids (Shen and Bargmann, 2003), and transgenic strains (injected at concentrations from 0.5-40ng/μl) using standard techniques. We coinjected relevant plasmids with markers *Punc-122::gfp*, *Punc-122::dsRed*, *Podr-1::gfp*, or *Podr-1::rfp* (15-40ng/μl) and generated the following transgenic strains: *olaEx866* [Pttx-3::snb-1::yfp, Pttx-3::mCh::rab-3], *olaIs10* [Pttx-3::gfp::syd-1, Pttx-3::mCh::rab-3], *olaEx1247* [Pttx-3::atg-9::gfp, Pttx-3::mCh::rab-3], *olaEx2266* [Pttx-3::mCh::rab-3], *olaEx2263* [Pttx-3::mCh::rab-3, Punc-14::atg-9::gfp], *olaEx2091* [Pttx-3::rfp, Pglr-3::rfp, Fosmid WRM0617aE11], *olaEx2169* [Pttx-3::rfp, Pglr-3::rfp, Patg-2::atg-2], *olaEx2281* [Punc-14::atg-9, Pttx-3::mCh, Pglr-3::mCh], *olaEx2188* [Pttx-3::rfp, Pglr-3::rfp, Fosmid WRM0616aF03], *olaEx2119* [Punc-14::lgg-1], olaEx2113[Patg-2::atg-2], *olaEx294* [Pttx-3::UtrCH::gfp], *olaEx1878* [Pttx-3::gfp::lgg-1, Pttx-3::mCh], *olaEx2558* [Pttx-3::gfp::lgg-1(G116A), Pttx-3::mCh], *olaEx2264* [Punc-14::atg-9::gfp, Pttx-3::mCh::rab-3], *olaEx2226* [Pdes-2::mCh], *olaEx2233* [Pdes-2::mCh::rab-3, Pdes-2::atg-9::gfp], *olaEx2221* [Patg-9(1.9kB upstream)::atg-9 genomic (first 6.4kB)::SL2::gfp, Pttx-3::mCh], *olaEx2191* [patg-2::gfp, Pttx-3::mCh]. Comprehensive subcloning information is available upon request.

We amplified ATG-9 and LGG-1 cDNA by PCR from a pool of cDNA from a mixed-stage population of animals.

We used a CRISPR protocol (Dickinson et al., 2015) to create *atg-9(ola270[atg-9::gfp::SEC])*, in which the eGFP coding sequence and the self-excision cassette are inserted in place of the *atg-9* stop codon.

### Fluorescence microscopy and confocal imaging

We used 40x Plan Fluor, NA 1.3, and 60x CFI Plan Apo VC, NA 1.4, oil objectives on an UltraView VoX spinning disc confocal microscope on a NikonTi-E stand (PerkinElmer) with a Hammamatsu C9100-50 camera to image fluorescently tagged fusion proteins (eGFP, GFP, YFP, RFP, and mCherry with excitation wavelengths of 488 or 561 nm) and transmitted light in live *C. elegans* at room temperature. We used Volocity software (Improvision by Perkin Elmer) to acquire and process images, using “Extended Focus” to show images as maximal projections. Additional processing of images, such as rotations and cropping, was conducted with Adobe Photoshop CS4, and figures were assembled with Adobe Illustrator CS4 (Adobe Systems Incorporated). We performed all quantifications on maximal projections of raw data. We immobilized worms using 10mM levamisole (Sigma) and oriented anterior to the left and dorsal up for all images.

### Mosaic Analysis

We performed mosaic analyses on *atg-9(bp564)*, *lgg-1(bp500)* and *atg-2(bp576)* mutant animals by expressing unstable transgenes with an appropriate rescuing construct and cytoplasmic markers in RIA and AIY (*olaex2281, olaEx2091* and *olaEx2169)* as previously described (Colon-Ramos et al., 2007; Herman, 2007; Yochem and Herman, 2003). We used a Leica DM5000 B microscope to analyze mutant animals for presence of the transgene as defined by presence of cytoplasmic RFP in AIY and/or RIA. For each condition, we scored the percentage of worms with rescued AIY presynaptic pattern.

### SNP Mapping and Whole-Genome Sequencing

We isolated *atg-9(wy56)* from visual forward-genetic ethyl methanesulfonate (EMS) mutagenesis screens designed to identify mutants that displayed abnormal AIY synaptic vesicle patterning (visualized with GFP::RAB-3). We used single-nucleotide polymorphism (SNP) mapping as described (Davis and Hammarlund, 2006; Davis et al., 2005) to map *wy56* to an interval between 0.005 Mb and 0.5 Mb on chromosome V.

We then performed whole-genome sequencing at the Yale Center for Genome Analysis (YCGA) on *wy56* animals as described (Bigelow et al., 2009; Sarin et al., 2008). We analyzed the data using MaqGene software and confirmed lesions by Sanger sequencing.

### AIY and PVD Quantifications

To quantify the penetrance of the AIY presynaptic defect, we used the integrated transgenic line *wyIs45* in the specified mutant backgrounds. Zone 2 was defined morphologically as the region of the AIY process which turns dorsally from the anterior ventral nerve cord into the nerve ring in adult animals (approximately 5 μm in length, and highlighted with a dashed box in all AIY figures). For a more detailed description of the quantification of penetrance, please see (Colon-Ramos et al., 2007; Stavoe et al., 2012; Stavoe and Colón-Ramos, 2012). We defined Zone 3 as the region of the AIY process dorsal to Zone 2 that extends to the end of the dorsal midline. These anatomical definitions were based upon EM reconstruction micrographs (White et al., 1986). We then scored the number of animals displaying normal or abnormal Zone 2 synaptic patterns relative to wild type animals.

To quantify the penetrance of F-actin enrichment, we used the stable extrachromosomal transgenic line *olaEx294* (Stavoe and Colón-Ramos, 2012). We scored the number of animals displaying a lack of enrichment of UtrCH::GFP in AIY Zone 2 relative to wild type animals as described (Stavoe and Colón-Ramos, 2012).

To quantify mutant animals in the L1 larval stage, we performed an egg lay as described (Porta-de-la-Riva et al., 2012) and examined resulting progeny after 14-18 hours for AIY presynaptic defects.

We quantified rescue of AIY presynaptic assembly in autophagy mutants by scoring the number of animals displaying a wild type localization pattern of GFP::RAB-3 or mCh::RAB-3 in AIY Zone 2 as described (Colón-Ramos et al., 2007; Stavoe et al., 2012; Stavoe and Colon-Ramos, 2012).

To quantify autophagosome (GFP::LGG-1) puncta, we used the stable extrachromosomal array *olaex1878* [Pttx-3::gfp::lgg-1, Pttx-3::mCh] and scored for presence or absence of LGG-1 puncta in the AIY neurite. We only scored animals where GFP signal could be detected in AIY. In wild type worms (n=169), 73% of animals had GFP::LGG-1 signal in AIY. We also note that although the same extrachromosomal transgenic line was used in different mutant backgrounds, GFP::LGG-1 signal was more prevalent in autophagy mutants as compared to wild type animals, with signal observed in 99% of *atg-3(bp412)* mutants, 100% of *atg-9(bp564)* mutants, 89% of *atg-2(bp576)* mutants, 100% of *epg-6(bp424)* mutants, 99% of *epg-5(tm3425)* mutants, 86% of *atg-2(bp576);unc-104(e1265)* double mutants, and 98% of *epg-6(bp424);unc-104* double mutants, (n>100 for all groups). We hypothesize that the absence of signal in 27% of wild type worms represents active autophagy degrading the LGG-1 substrate. While we note these observations, they do not affect the interpretations presented in the manuscript.

We measured the PVD axon length along the ventral nerve cord (which does not include the distance between the cell body and the ventral nerve cord) using ImageJ software.

### Statistical Analyses

To determine statistical significance for categorical data, we used Fischer’s exact test. Error bars represent 95% confidence intervals.

Statistical significance for continuous data was determined using one-way ANOVA with *post hoc* analysis by Tukey’s multiple comparisons test using PRISM software. Error bars for continuous data were calculated using standard errors of the mean (s.e.m.).

## AUTHOR CONTRIBUTIONS

AKHS, SEH and DACR designed the experiments; AKHS and SEH performed the experiments and data analyses. AKHS, SEH and DACR prepared the manuscript.

## ACKNOWLEDGEMENTS

We thank members of the Colón-Ramos lab and Erika Holzbaur (University of Pennsylvania) for their thoughtful comments on the manuscript and the project. We are particularly indebted to Kang Shen (Stanford University) for reagents, thoughtful comments and advice during the course of this project. We thank summer interns Jihane Jadi (University of North Carolina, Chapel Hill) and Ninoshka Caballero-Colón (University of Puerto Rico, Humacao) for experimental assistance. We thank the *Caenorhabditis* Genetics Center for mutant strains, Cori Bargmann (Rockefeller University) for CX9797 (kyIs445) strain, David Miller (Vanderbilt University) for NC1686 (wdIs51) strain, the Mitani laboratory of the Tokyo Women’s Medical University School of Medicine for autophagy mutant strains, and the Hong Zhang laboratory at the Institute of Biophysics, Chinese Academy of Sciences for autophagy mutant alleles. We thank Z. Altun (www.wormatlas.org) for diagrams used in figures. We thank Bob Goldstein (University of North Carolina at Chapel Hill) for CRISPR Self-excision cassette (SEC) constructs and advice.

We thank the Research Center for Minority Institutions program and the Instituto de Neurobiologla de la Universidad de Puerto Rico for providing a meeting and brainstorming platform. This work was partially conducted at the Marine Biological Laboratories at Woods Hole, under a Whitman research award to DACR. This work was funded by the U01HD075602, R01NS076558 and R24OD016474 NIH grants to DACR. AKHS and SEH were supported by Cellular and Molecular Biology Training grant T32-GM007223 from the NIH. SEH was also supported by NSF Graduate Research Fellowship DGE-1122492.

